# Characterization of Novel Recombinant Mycobacteriophages derived from Homologous Recombination between two Temperate Phages

**DOI:** 10.1101/2022.12.15.520664

**Authors:** Hamidu T Mohammed, Catherine Mageeney, Jamie Korenberg, Lee Graham, Vassie C Ware

## Abstract

Comparative analyses of mycobacteriophage genomes reveals extensive genetic diversity in genome organization and gene content, contributing to widespread mosaicism. We previously reported that the prophage of mycobacteriophage Butters (cluster N) provides defense against infection by Island3 (subcluster I1). To explore the anti-Island3 defense mechanism, we attempted to isolate Island3 defense escape mutants on a Butters lysogen, but only uncovered phages with recombinant genomes comprised of regions of Butters and Island3 arranged from left arm to right arm as Butters-Island3-Butters (BIBs). Recombination occurs within two distinct homologous regions that encompass *lysin A, lysin B*, and *holin* genes in one segment, and *RecE* and *RecT* genes in the other. Structural genes of mosaic BIB genomes are contributed by Butters while the immunity cassette is derived from Island3. Consequently, BIBs are morphologically identical to Butters (as shown by transmission electron microscopy) but are homoimmune with Island3. A reverse experiment where an Island3 lysogen was infected with Butters yielded Butters phages and no recombinants, demonstrating directionality to the recombination phenomenon. Recombinant phages overcome antiphage defense and silencing of the lytic cycle. We leverage this observation to propose a stratagem to generate novel phages for therapeutic use.

**GRAPHICAL ABSTRACT:** 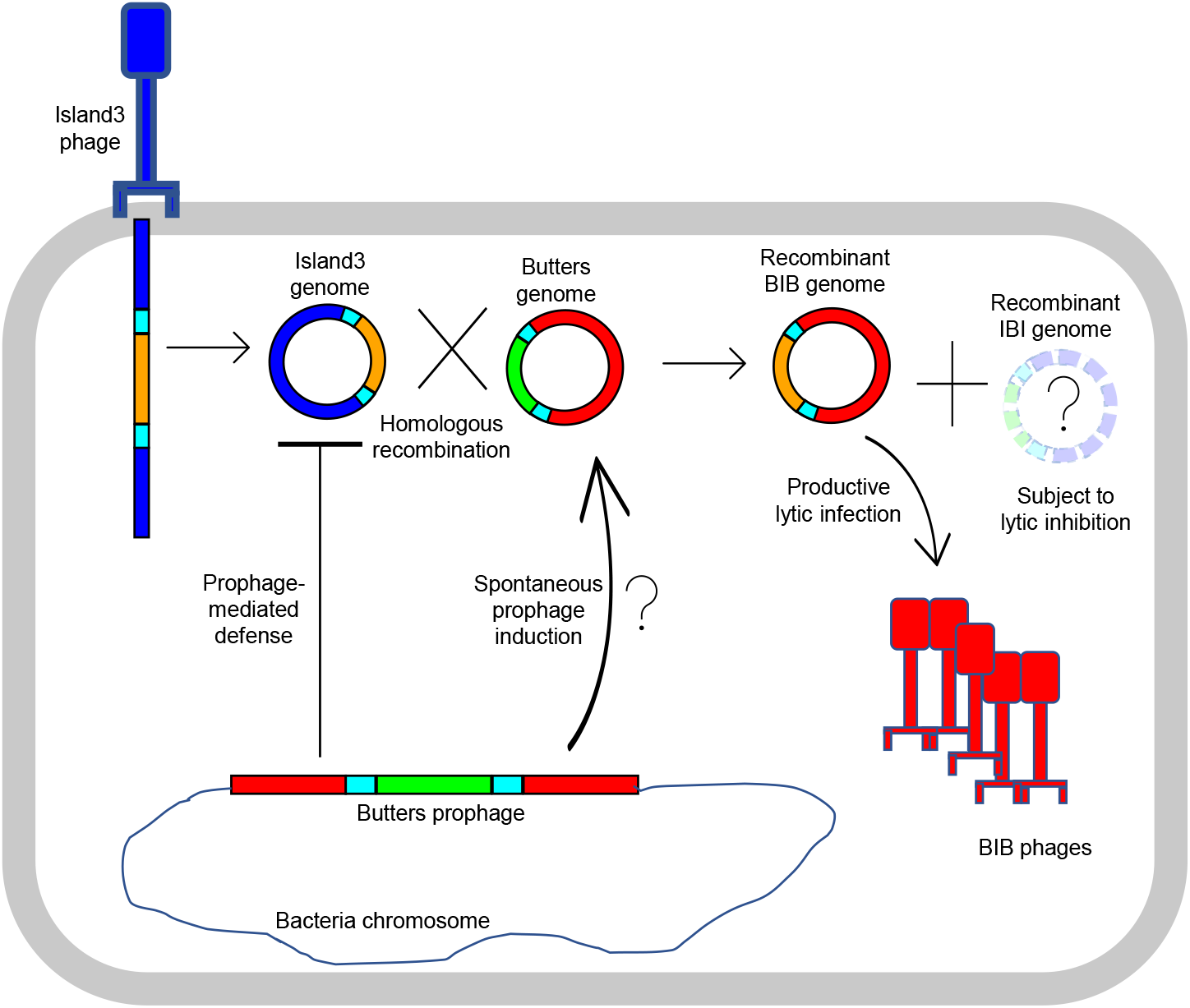

## INTRODUCTION

The uptick in phage biology research coupled with a more accessible sequencing regime has led to sequencing thousands of phage genomes (Reyes *et al*. 2012). Comparative genome analysis reveals extensive mosaicism that characterizes phage genomes (Mavrich and Hatfull 2017; Oliveira *et al*. 2019; Yahara *et al*. 2019). Phage genomes are conceived of as a patchwork of exchangeable genetic modules that are shuffled between phages via horizontal gene transfer (Botstein). Phage genome mosaicism is generated mechanistically by homologous recombination (Clark *et al*. 2001; Martinsohn *et al*. 2008; De Paepe *et al*. 2014) or by illegitimate recombination (Hendrix *et al*. 2000; Juhala *et al*. 2000; Pedulla *et al*. 2003; Morris *et al*. 2008). With either recombination mechanism, the opportunity for recombination between phage genomes would be presented in a host cell during coinfection or when an invading phage infects a lysogen (Hendrix *et al*. 2000).

The prevalence of prophages in bacterial genomes (Casjens 2003; Touchon *et al*. 2016) likely increases the probability of an invading phage genome co-occurring along with a resident prophage, creating an avenue for recombination between two phage genomes. Some prophages collude with their host to defend against heterotypic phage infection (Labrie *et al*. 2010; Bondy-Denomy *et al*. 2016; Dedrick *et al*. 2017; Hampton *et al*. 2020). It is theorized that genetic recombination between phages may be an evolutionary strategy for phages to exchange antiphage defense systems (Kauffman *et al*.; Piel *et al*. 2021).

The availability of large collections of sequenced mycobacteriophages provides opportunity to observe and investigate the relationship between phage genome mosaicism and exchange of antiphage defense systems. The HHMI-sponsored SEA-PHAGES program (“The Actinobacteriophage Database | Home”; Russell and Hatfull 2017; Hatfull 2020), in which over 3,000 phages of the >20,000 infecting Actinobacteria have been sequenced, provides such a reservoir. The majority (ca. 12,242) of those sequenced phages are mycobacteriophages isolated on the soil organism *Mycobacterium smegmatis* mc^2^155 (“The Actinobacteriophage Database | Home”). These genomes are extensively mosaic (Pedulla *et al*. 2003; Pope *et al*. 2015). Mycobacteriophages are currently grouped into 31 clusters (A-Z, AA-AE) based on genome similarity and a singletons with no known close relatives (“The Actinobacteriophage Database | Home”; Cresawn *et al*. 2011; Pope *et al*. 2015; Russell and Hatfull 2017). Although genomes are categorized into clusters, many genes and groups of genes are shared between clusters, thereby contributing to extensive genome mosaicism.

Cluster N mycobacteriophages, in particular, have been highlighted for the diversity that characterizes the central region of their genomes between more conserved sequences in the left and right arms (Dedrick *et al*. 2017). Genes within the central region of cluster N mycobacteriophages are expressed in cluster N lysogens and have been implicated in prophage-mediated defense mechanisms that defend the host from heterotypic viral attack (Dedrick *et al*. 2017; Mageeney *et al*. 2020). Cluster N and I1 mycobacteriophage genomes are notable in their continuum of genetic similarity and are proposed to exchange genetic modules at a relatively faster rate (Mavrich and Hatfull 2017). Despite extensive *in silico* characterization of mosaicism within mycobacteriophage genomes (Pedulla *et al*. 2003; Morris *et al*. 2008; Dedrick *et al*. 2017; Mavrich and Hatfull 2017), relatively little direct experimental evidence exists to document how mosaicism emerges. Thus, gaps remain in our understanding of mechanisms underlying development of genome mosaicism and in the biological consequences of this phenomenon among mycobacteriophages.

Here, we describe a homologous recombination event between the Butters (cluster N) prophage and the Island3 (subcluster I1) genome that yields a novel mosaic recombinant phage named BIB (Butters-Island3-Butters). Recombinants form when Butters acts as the prophage and Island3 acts as the invading phage, but not vice versa. Our data reveal a biological phenomenon where homologous recombination provides a means for the invading phage to overcome antiphage defense and the resident phage to escape from lytic repression. We discuss this phenomenon in the context of phage therapy as a measure where mixed phage lysates in co-infections are used to control bacterial infections.

## MATERIAL AND METHODS

### Phage isolation, propagation, and genomic analysis

Phages (GenBank accession numbers KC576783 [Butters], HM152765 [Island3], and MN945904 [Eponine]) were isolated and grown on *Mycobacterium smegmatis* mc^2^155 as described in (Caratenuto *et al*. 2019) Island3 lysate was obtained from the Hatfull lab (University of Pittsburgh). The genomic sequence for the Island3 strain used in this study differs from that of the wild type with a 257-bp deletion (coordinates 43307 to 43563) and a C2656T SNP. DNA for recombinant BIBs were isolated using phenol:chloroform extraction. BIBs 1-4 were sequenced at the Pittsburgh Bacteriophage Institute, Pittsburgh, PA. and BIBs 6-10 were sequenced at Novogene Corporation Inc. Phage lysates (titers, **≥**1×10^9^ PFU/ml), diluted with phage buffer (0.01 M Tris, pH 7.5, 0.01 M MgSO_4_, 0.068 M NaCl, and 1 mM CaCl_2_), were used for immunity testing and PCR. Phamerator (Cresawn *et al*. 2011) was used for comparative genomic analysis and genome map representation.

### Phage Sensitivity Assays

*Spot test assay for immunity tests:* Lawns of *M. smegmatis or* lysogens were made by plating 250 μl of the bacteria culture with 3.5 ml of top agar (at ∼55°C) on an LB agar plate (with 10 μg/ml of cycloheximide [CHX] and 50 μg/ml of carbenicillin [CB]). Phage lysates were serially diluted to 10^−7^ and spotted (3μl each) onto lawns of interest. Plates were incubated for 48 h at 37°C. Phage growth was assessed at 24 and 48 h. *Plaque assays:* Phage lysates were serially diluted. 10 μl of each dilution was added to 250 μl of the culture or lysogen to be infected and incubated for 10 minutes at room temperature. Top agar (3.5 ml) was added to the phage-bacteria mix and plated on LB agar plates (CHX/CB).

### Electron Microscopy

Phage lysate was deposited onto 200 mesh copper grids coated with formvar and carbon (Ted Pella: 01810), rinsed with deionized water and stained with 2% ammonium molybdate. Samples were imaged on a JEOL 1200EX transmission electron microscope. Capsid and tail length measurements were done using ImageJ (Schneider *et al*. 2012). Measurements represent the average of at least 6 particles.

### Polymerase Chain Reactions

Samples were derived from pure phage lysates or plaques picked into 100 μl of phage buffer. Platinum™ II Hot-Start Green PCR Master Mix (2X) (Invitrogen) was used. 1 μl of each sample was used as template. Primers used are listed in Table 1. All primers were ordered from Integrated DNA Technologies, Inc.

### Determination of number of secondary plaques to screen

Assuming 1 out of 10 phages within a plaque is non-BIB (i.e., either Butters, Island3, or the reciprocal product: Island3-Butters-Island3 [IBI]), then the probability of obtaining 1 non-BIB phage at 95% confidence interval is given by:

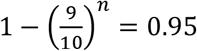, where n is the sample size of plaques to be screened to achieve a 95% confidence interval. From this equation we calculated that *n* must be at least 28.5 (rounded up to 30).

## RESULTS

Mycobacteriophage Butters is a temperate phage that encodes distinct antiphage defense systems. Its repressor blocks infection by other cluster N phages (Dedrick *et al*. 2017). Additionally, Butters blocks infection by the heterotypic phage Island3 through a repressor-independent system (Dedrick *et al*. 2017; Mageeney *et al*. 2020). To explore the mechanism of defense and to identify putative Island3 targets of Butters prophage defense, we pursued isolation of Island3 defense escape mutants (DEMs) - mutants that have escaped Butters prophage-mediated defense and consequently, can efficiently infect a Butters lysogen [mc^2^155(Butters)].

Single Island3 plaques, which typically constitute evidence of mutants, were not observed when serial dilutions of an Island3 high titer lysate were spotted on a Butters lysogen lawn (Figure 1). Prominent spot clearings of high Island3 concentrations are regarded as “killing from without” rather than evidence of successful infections (Abedon; Delbrt3ck). Yet, it is conceivable that clearings associated with “killing from without” may mask individual plaques arising from an Island3 DEM. To explore this possibility, plaque assays were performed.

**Figure 1.**
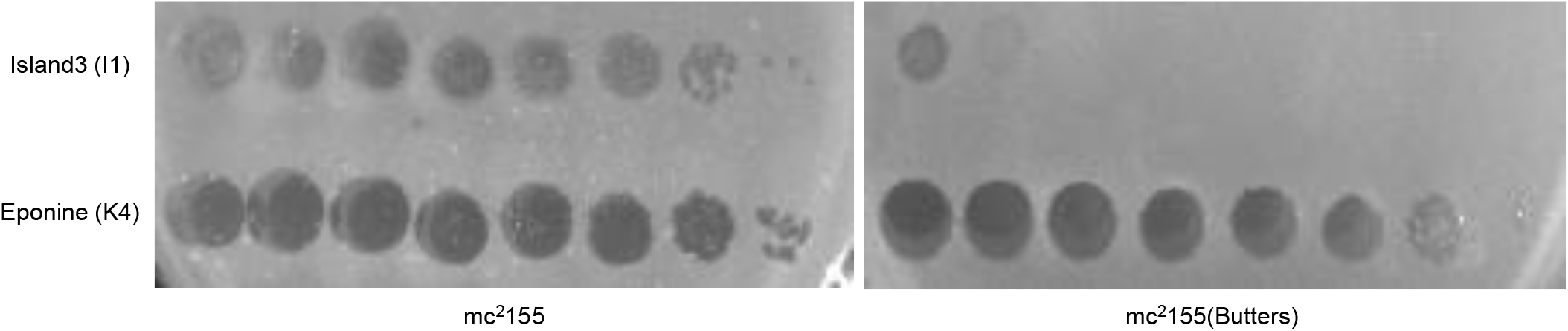
Phage sensitivity assay (spot tests) showing mc^2^155 (Butters) defense against Island3. Ten-fold serial dilutions (3 μl) of Island3 and Eponine (as control) were spotted onto lawns of *Mycobacterium smegmatis* mc^2^155 (mc^2^155) and Butters lysogen [mc^2^155(Butters)].

Typically, about ten plaques are formed upon infecting Butters lysogen with 10^8^ pfu of Island3. Nine of these putative Island3 DEMs were successfully sequenced.

### Recombinant plaques arise from a double homologous recombination event between Butters and Island3 genomes

Sequencing of putative Island3 DEMs showed that genomes are recombinants with the left arm contributed by Butters, the middle segment derived from Island3, and the right arm contributed by Butters (Figure 2A). These recombinant phages are termed **B**utters-**I**sland3-**B**utters (BIBs). Comparative genome analysis reveals that BIBs are a product of a double homologous recombination event within two highly homologous regions between Butters and Island3. The first homologous region has an average nucleotide identity (ANI) of about 84% and spans about 2727bp of both genomes (Figure 2A). This homology region contains four predicted open reading frames (ORFs) or protein coding genes; lysin A, lysin B, holin, and a gene of unknown function (Butters gene *29* or Island3 gene *32*) (Figure 2A). By mapping single nucleotide polymorphisms (SNPs) originating from Butters versus those originating from Island3 to BIB genome sequences, we established that the crossover switch from the Butters genome to Island3 genome occurs within lysin B (Figure 2B). There are instances where genetic exchange occurs at several positions, as revealed by the presence of multiple SNPs from either phage genome occurring within the genomic region of interest. The second homology region has an ANI of about 98% and spans about 2237bp of both genomes (Figure 2A). This second homology region contains three ORFs: a gene of unknown function (Butters gene *49* or Island3 gene *47*), RecE, and RecT (Figure 2A). Here, the exact crossover point varies within RecE and RecT (Figure 2A). Evidence for multiple points of genetic exchange is shown by the presence of alternating Butters or Island3 SNPs among different BIBs (Figure 2C).

**Figure 2:**
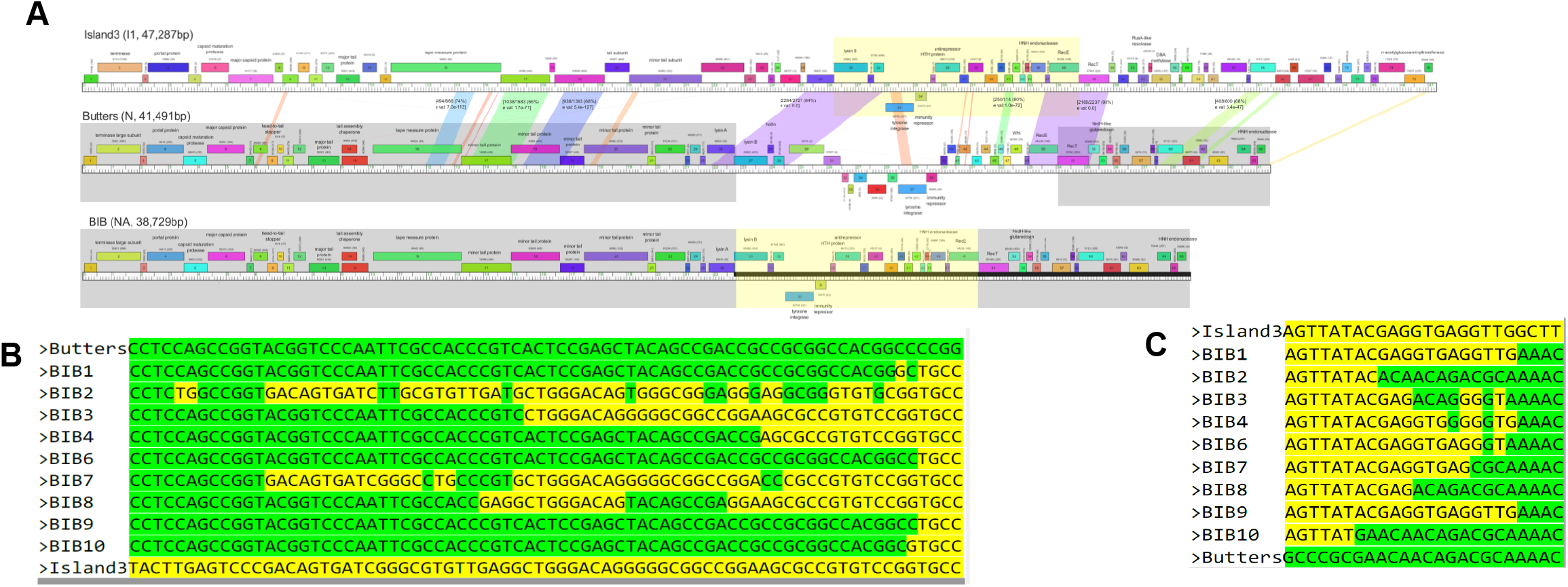
Genome characterization of Island3, Butters, and recombinant phage, BIB. **(A)** Phamerator maps (Cresawn *et al*. 2011) for Butters, and Island3 and a derived map for BIB. Genes are depicted by colored rectangular boxes and are aligned to their genomic ruler.Gene numbers are within the rectangular boxes and putative gene functions, if known, are indicated above the gene. Numbers above the genes represent gene group names or phams and numbers in parenthesis show number of genes within the pham. Color-coded patches connecting genomes show nucleotide similarity between the genome segments with red being the least similar (E value = 1e-4) and violet being the most similar (E value = 0). Selected segments with over 300bp aligned sequences and E value ≤ 1e-20 are shown within the color-coded connecting patch. Within each square bracket is the fraction of number of nucleotide identities over total aligned sequences, percentage of nucleotide identity in parentheses and E value (e val). The bottom row is a representative map of a BIB genome. Blocks or modules derived from Butters are shaded transparent dark and module derived from Island3 is shaded transparent yellow. **(B)** Alignment of SNPs across the first homology region. The entire nucleotide sequences that defines the first homology region (genome coordinates: 21825-23917 [Butters] and 25302-27409 [Island3]) were aligned in Jalview (Waterhouse *et al*. 2009). Conserved sequences were redacted to yield only SNPs. From left to right, sequences of BIBs first map to Butters before transitioning into Island3. SNPs in some BIBs “flipflop” from one parental genome to another and back. **(C)** Alignment of SNPs across the second homology region (genome coordinates: 33050-35516 [Butters] and 33774-35854 [Island3]. From left to right, BIB genomes align to Island3 before transitioning into Butters.

### Recombinant BIB phenotypic characteristics are determined by specific parental phage genomic modules

Morphological and immunity characteristics of recombinant BIBs were compared to their parental phages. Plaque morphologies of mycobacteriophages differ on different media (unreported data). Plaque morphologies described here were determined on 7H10 agar. After ∼36-48 hours on incubation, Butters forms 2.0 - 3.0 mm turbid plaques with a translucent center encircled by a cloudier ring (Figure 3A). Island3, on the other hand, forms 2.0 - 2.5 mm diameter bullseye plaques with three distinct regions: a clear center, a cloudy ring enveloping the clear center, and a clear outer ring (Figure 3A). Interestingly, BIBs show an emergent plaque morphology, distinguishable from those of Butters or Island3. While BIBs form bullseye plaque morphologies like those of Island3, the clear centers of BIB plaques are relatively larger while the middle cloudy rings are significantly smaller (Figure 3A). It therefore appears that genes within the exchanged module from Island3 play a significant part in determining plaque morphology phenotype; however, contributions from the non-Island3 part of the BIB genome cannot be ruled out.

**Figure 3.**
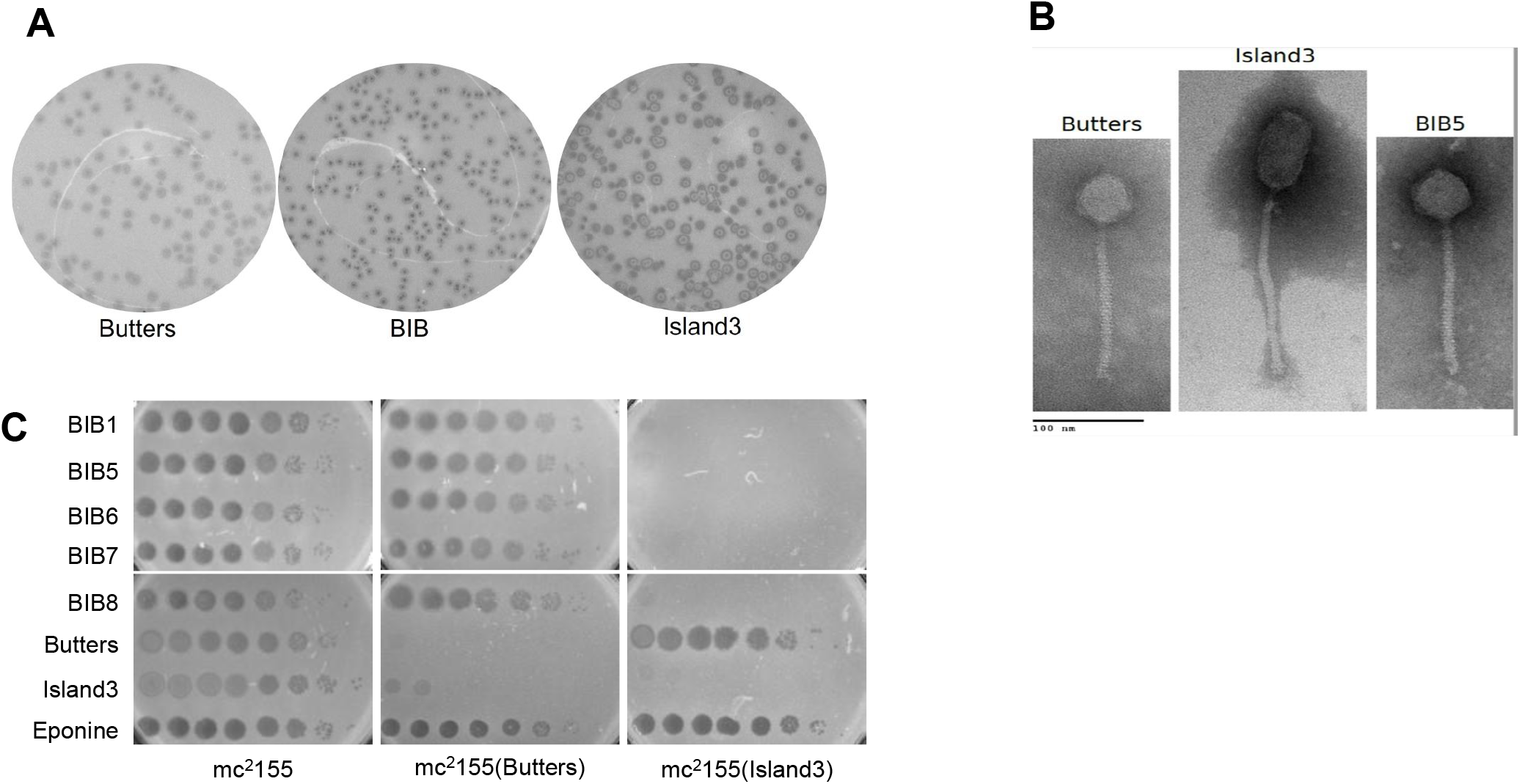
Physical and immunity properties of BIBs and their parental phages. **(A**) Comparison of plaque morphologies of Butters, Island3 and BIBs. Plaque assay was performed by plating serial dilutions of the phage lysates onto mc^2^155 and plaque morphologies assessed. BIBs present an emergent phenotype distinguishable from that of both Butters and Island3. **(B)** Electron micrograph of Butters, Island3 and BIB phages. BIBs and Island3 share identical capsid and tail lengths. **(C**) Immunity relationships for Butters, Island3 and BIBs. 3 μl of serially diluted phages were spotted on lawns of mc^2^155, mc^2^155(Butters) and mc^2^155(Island3). BIBs are homoimmune with Island3.

Given that BIBs derive all structural assembly genes from Butters and the immunity cassette from Island3, we hypothesized that BIBs will be morphologically identical to Butters and homoimmune with Island3. Transmission Electron Microscope (TEM) images show both BIBs and Butters have an icosahedral head with a diameter of 53.0 ± 4.0 nm while Island3 has a prolate head with longitudinal height of 82.0 ± 2.0 nm and latitudinal width of 41.0 ± 2.0 nm (Figure 3B). Further, both BIBs and Butters have tail lengths of 169.0 ± 5.0 nm while that of Island3 is 202.0 ± 2.0 nm (Figure 3B). These results are not surprising, as both major capsid protein (which defines capsid diameter) and tape measure protein (which define tail lengths) of BIBs are contributed by Butters. Immunity experiments performed by spotting dilutions of BIB lysates on lawns of *M. smegmatis* mc^2^155, mc^2^155(Butters) and mc^2^155(Island3) show that BIBs efficiently infect mc^2^155(Butters), but are inhibited on mc^2^155(Island3) lawn, due to Island3 immunity repressor activity (Figure 3C).

### BIB plaques are the predominant type from Island3 infection of mc^2^155(Butters)

Since 100% of all plaques isolated and sequenced were recombinant BIBs, we investigated whether plaques produced from Island3 infection of mc^2^155(Butters) were of mixed types that included wild type forms of Island3 and Butters and the reciprocal recombinant product IBI. Plaque assays were performed using Island3 to infect mc^2^155(Butters) to generate primary plaques. Primers were designed that would specifically PCR amplify each phage type across the recombinant regions. All primary plaques showed a BIB-specific product but were negative for the other predicted phage types (Figure 4A, data shown for PCR screening across first homology region; PCR data for second). Additionally, genome sequencing confirmed exchange between both homology regions (see BIB GenBank accession numbers provided in Methods). To test the possibility that the experimental set up may fail to detect other phage types present at low concentration, a null hypothesis was established, predicting that there would be a ratio of at least one phage type of Island3, IBI, or Butters to nine BIBs within a primary plaque. If the hypothesis is supported, screening 30 secondary plaques resulting from a plating a primary plaque on mc^2^155 should yield at least one Island3, IBI, or Butters plaque at a 95% confidence interval (see methods). Plaque assays for three primary plaques were performed to generate secondary plaques on mc^2^155. In all cases, all thirty plaques were positive for a BIB-specific band but negative for all other phage types [Supplementary Figure S1 (data for one set shown)]. The lack of other phage types essentially rules out the possibility of amplification of the Island3 genome prior to recombination as well as any amplification of an IBI genome, if formed.

**Figure 4.**
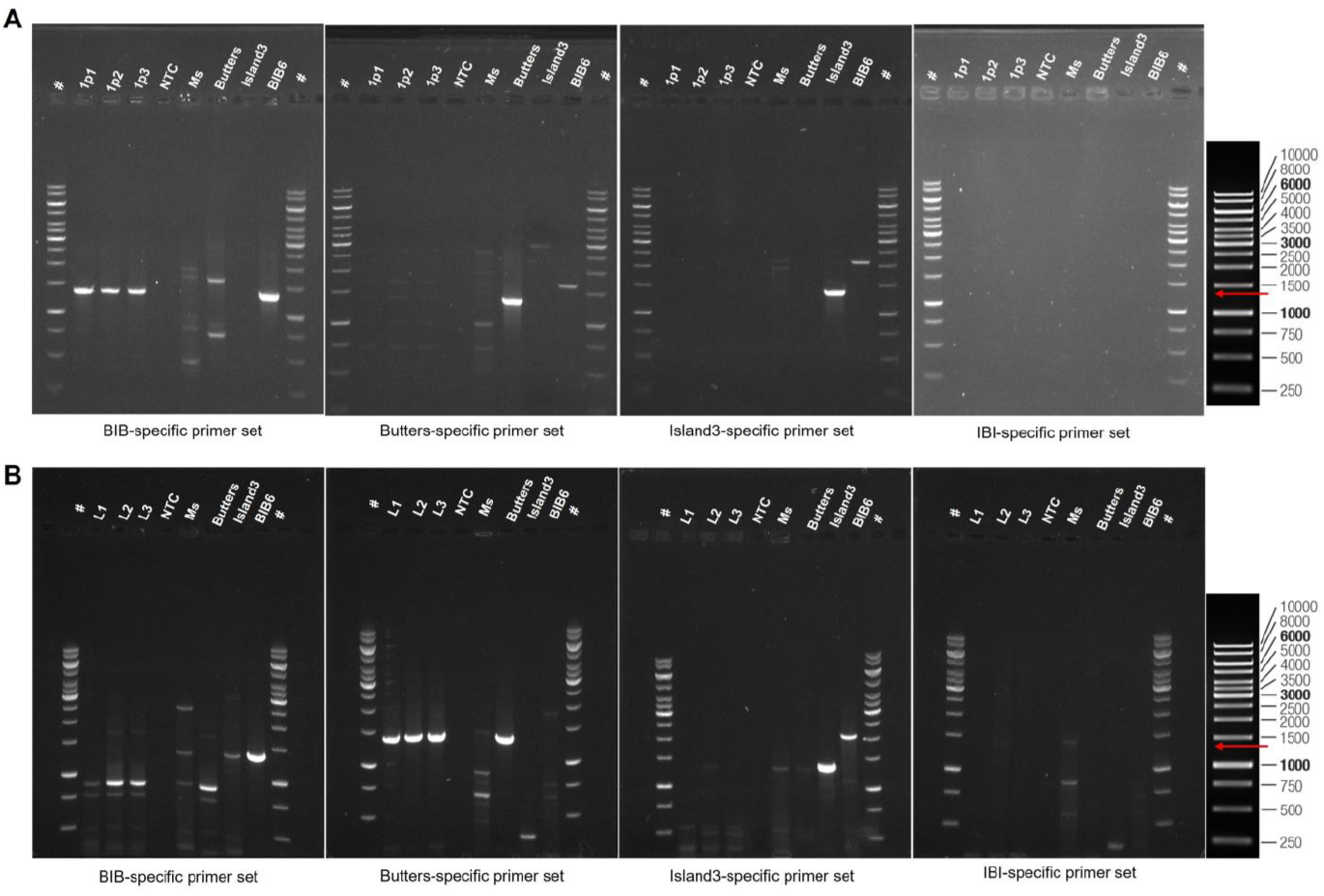
PCR screening of plaques obtained from plating experiments. Primers specific for each possible phage type (BIB, Butters, Island3, and IBI) for first homology region were designed and used to screen plaques via PCR. PCR products were run on 0.8% agarose gel at 60V. GeneRuler 1kb DNA ladder, ThermoFisher Scientific (extreme right panel) was used as ladder (#). Expected product size is 1367bp for BIB (red arrow). Non-specific bands appear in some instances due to non-specific primer annealing for PCR conditions used. Also, some of the non-specific bands can be mapped to mc^2^155 genome which is inevitably present when crude phage lysates are filtered through 0.2 μm filters. **(A)** PCR screening of plaques obtained from plating Island3 on mc^2^155(Butters). Samples from left to right are ladder, primary plaques (1p1, 1p2 and 1p3), no template control (NTC), *M. smegmatis* mc^2^155 *(Ms)*, Butters lysate, Island3 lysate, BIB6 lysate, and ladder (#). Note that primary plaques show PCR products with BIB-specific primers but not with Butters-or Island3-or IBI-specific primers. **(B)** PCR screening of lysates obtained from plating Butters lysate on mc^2^155(island3). Samples from left to right are ladder, lysates (L1, L2 and L3), no template control (NTC), *M. smegmatis* mc^2^155 *(Ms)*, Butters lysate, Island3 lysate, BIB6 lysate, and ladder (#). Note that in the reciprocal experiment, products are detected only with Butters-specific primers (expected product size is 1361bp).

### Recombinants form when Butters, but not Island3, is the prophage

To test whether IBI recombinant phages are produced when Island3 is the prophage, reciprocal experiments to plate a Butters lysate on mc^2^155(Island3) were performed. No IBI-specific or BIB-specific PCR products were detected from plaques and lysates from these experiments (Figure 4B). All samples yielded Butters-specific PCR products only (Figure 4B). These results establish directionality to the observed recombination phenomenon.

## DISCUSSION

The phenomenon of extensive mosaicism is well documented within phage genomes in general and within mycobacteriophages in particular (Mavrich and Hatfull 2017; Oliveira *et al*. 2019; Yahara *et al*. 2019). Evidence supporting mosaicism within mycobacteriophage genomes is derived largely from bioinformatic analyses of published genomes. While informative, these post-facto analyses reveal little, if any, information about factors and evolutionary pressures that trigger or limit the occurrence of recombination between phage genomes that lead to mosaicism. Here, wet-lab evidence shows homologous recombination between two temperate mycobacteriophages, thus providing the springboard for a more mechanistic interrogation of how genomic mosaicism is generated within mycobacteriophages.

Comparative genome analysis suggests that the observed recombination is likely a double crossover event at two highly homologous regions between the Butters prophage and Island3 genomes. Butters and Island3 have 68% ANI. While there are several other homologous regions across the two genomes with ANI ranging from 43-80%, all sequenced BIBs recombined at the two regions with the highest ANI of 84% and 98%, respectively. It is unknown if recombinant phage form by homologous recombination at a single homologous region (to yield BI or IB) or by recombination at any other potential homologous regions, as no evidence of these possibilities was apparent. Some BIB genomes show evidence of multiple genetic exchanges (e.g., “flip-flop” of SNPs) between one parental genome to the other and then back. These “flip-flops” are likely gene conversion events resulting from resolution of Holliday junctions (Santoyo and Romero 2005; Guirouilh-Barbat *et al*. 2014).

Structural assembly genes determine phage morphotype (Katsura and Hendrix 1984). Expectedly, EM morphology of BIBs is similar to that of Butters with a comparable tail length of about 169.0 ± 5.0 nm and head diameter of about 53.0 ± 4.0 nm (Figure 3B). Island3, on the other hand presented a longer tail length of about 202 ± 2.0 nm and a prolate head with a longitudinal height of about 82.0 ± 2.0 nm and latitudinal width of 41.0 ± 2.0 nm (Figure 3B). Homoimmunity of BIBs with Island3 was also expected since the BIB immunity cassette is derived from Island3. Several factors affect plaque morphology including lysogeny (Levine 1957; Shin *et al*. 2014)), the presence of depolymerases on tail proteins (Pires *et al*. 2016), phage tolerance (Tzipilevich *et al*. 2021) and variation of media or growth conditions (Ramesh *et al*. 2019). All experiments were carried out on 7H10 plates using the same batch of mc^2^155 for each biological replicate. BIBs plaques are morphologically similar to those of Island3, yielding bullseye plaques in both instances, but the defining concentric circles are markedly different in areas. Clearly, the exchanged genetic module derived from Island3 contributes genes that significantly affect plaque morphology phenotype. Yet, the inability to fully recapitulate the Island3 plaque morphology in BIBs suggests that other genes native to Butters portions of BIBs play a role in determining plaque morphology.

Evolutionary pressures and other biological phenomena that drive recombination between phage genomes are poorly understood. A recent study reported that the *Bacillus subtilis* prophage SPβ recombines with phi3T DNA during sporulation, but not in the absence of sporulation (Dragoš *et al*. 2021). It has been suggested that recombination is a means for phages to escape antiphage defenses (Kauffman *et al*.; Piel *et al*. 2021). In support of this, Piel *et al*. (2021) (Piel *et al*. 2021) showed that Vibrio phages escape antiphage defenses by exchanging anti-defense genes through homologous recombination (Piel *et al*. 2021). In this study, the outcome of recombination is that BIB phages escape both prophage-mediated defense against Island3 and repressor-mediated defense against Butters.

Recombinant IBIs were neither recovered from infections of Island3 on a Butters lysogen nor from the reciprocal experiment where Butters was plated on mc^2^155(Island3). In the former case, we would expect an IBI to be inhibited from the repressive action of Butters immunity repressor. Such repression should be absent in the latter case and hence we would expect IBIs to be propagated, if formed. Several reasons may account for the absence of IBIs. First, it is possible that IBIs are non-viable. Within the exchanged modules, Island3 genes *33 (integrase), 34 (repressor), 35 (HTH protein) and 36 (antirepressor)* are annotated to function in lysis-lysogeny regulation. These genes are replaced by Butters genes *37 (integrase), 38 (repressor), and 39 (excise)* which modulate lysis-lysogeny outcomes in Cluster N mycobacteriophages(Broussard *et al*. 2013). Island3 genes *37, 41, 47, and 48 (RecE)* have homologues within the Butters exchange module corresponding to Butters genes *40, 45, 49, and 50 (RecE)*, respectively. Crucially, Island3 genes *38-40, 42-45, 46 (*a type of *HNH endonuclease)* have no homologues within the Butters module. If any of these Island3 genes play essential roles within the lytic cycle of Island3, then absence of a functional Butters homologue within the Butters module would account for IBI non-viability.

Secondly, while recombination may be pervasive among groups of phages and lead to an increase in gene content variation/diversity, genome size remains conserved (Piel *et al*. 2021) presumably due to constraints imposed by capsid size. The Butters exchange module is ∼2700bp larger than the Island3 exchange module. This size increase would augment the genome of a hypothetical IBI to ∼50,000bp instead of 47000bp. It is unknown if the prolate head of a recombinant IBI phage could accommodate the increase in genome size. Thirdly, it is unknown which phage factors (from Island3 or Butters) drive recombination. Detection of only one recombinant type suggests a directionality to the recombination phenomenon and potentially unequal contributions of phage factors to facilitate the recombination event. Further experimentation will address this question.

This work parallels that of De Paepe *et al*. (2014) (De Paepe *et al*. 2014) where recombination between invading temperate phages and cryptic prophages in *E. coli* was observed(De Paepe *et al*. 2014). Their findings supported the proposition that recombinants arise after several rounds of phage genome amplification (Visconti and Delbrück 1953). Failure to detect Island3-specific PCR products within primary plaques is due to the presence of a prophage-mediated system which inhibits the generation of infective Island3 particles. The interplay between the Butters prophage-mediated defense system and the observed recombination is yet unknown. The proposal of Kauffman *et al*. (2021) (Kauffman *et al*.) that prophage-encoded antiphage defenses may digest genomes of invading phages, yielding fragments that recombine with the prophage, may be applicable in generating BIBs in this system.

### Targeted phage-phage recombination as a strategy to overcome dearth of therapeutic phages

The growing incidence of antibiotic resistance in bacterial pathogens has revived interest in phage therapy. Several cases of emergency use authorization of phages for treatment of drug-resistant infections have been reported with varying degrees of success (Kutateladze and Adamia 2008; Aslam *et al*. 2019; Law *et al*. 2019; Dedrick *et al*. 2019). While promising, it is challenging to obtain phages capable of infecting a given drug-resistant pathogenic bacterial strain (Dedrick *et al*. 2021; Rebekah M. Dedrick, Bailey E. Smith, Madison Cristinziano, Krista G. Freeman, Deborah Jacobs-Sera, Yvonne Belessis, A. Whitney Brown, Keira A. Cohen, Rebecca M. Davidson, David van Duin, Andrew Gainey, Cristina Berastegui Garcia, C.R. Robert George, Ghady 2022). Typically, determining which phages infect a given bacterial strain is accomplished by plating pure phage lysates on the host. Some bacteria may carry antiphage defense systems capable of blocking any two phages A and B; yet co-infection by phages A and B could result in recombination to yield phages that escape the antiphage defense systems (Figure 5). This strategy could expand the pool of therapeutic phages especially for pathogenic bacteria for which phages are scarce. For pathogenic bacteria harboring resident prophage(s), therapeutic phages might be engineered to elicit recombination with the resident prophage. Our proposed strategy could employ both lytic and temperate phages. Resulting temperate phages that infect a given bacterial candidate for phage therapy could be engineered to yield a lytic phage.

**Figure 5.**
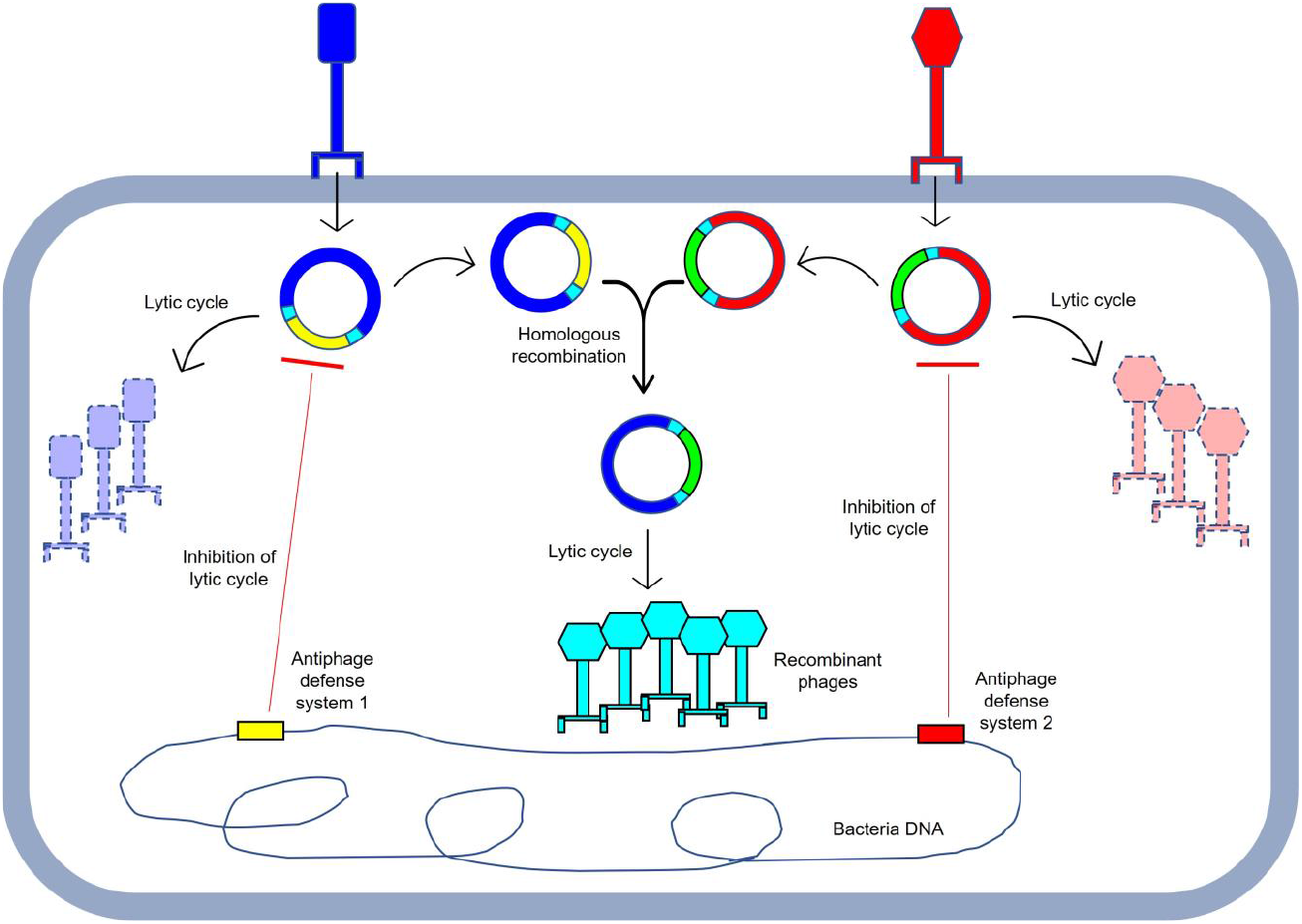
Model showing coinfection as a strategy to overcome antiphage systems. Bacteria (or lysogen) have antiphage defense systems that block lytic infection by both blue and red phages. Antiphage defense system 1 (yellow rectangular block) targets yellow portions of blue phage and antiphage system 2 (red rectangular block) targets red portions of red phage. During coinfection, homologous regions (cyan colored) may recombine, yielding recombinant phage that escape defense systems present in the cell.

This work provides the foundation for addressing several interesting questions about recombination within mycobacteriophages. (1) Does recombination occur before Butters prophage induction or is the prophage induced before recombination? (2) Are there any Butters prophage-or Island3-specific factors that trigger the observed homologous recombination? (3) What is the source of recombineering enzymes that catalyze the observed recombination? Potential recombineering candidates include the host mc^2^155 RecA, Butters *RecET* system or Island3 *RecET* system (Martinsohn *et al*. 2008; De Paepe *et al*. 2014). (4) Can recombination be reproduced within other mycobacteriophage clusters and subclusters? Overall, discovery of these naturally occurring mycobacteriophages that recombine to yield mosaic recombinant phages will allow interrogation of molecular mechanisms underpinning this phenomenon.

## DATA AVAILABILITY

GenBank accession numbers for phage genomes in this report are as follows: Butters (KC576783), Island3 (HM152765), Eponine (MN945904), BIB1 (OP961731), BIB2 (OP961730), BIB3 (OP961729), BIB4 (OP961728), BIB6 (OP961727), BIB7 (OP961726), BIB8 (OP961725), BIB9 (OP961724), and BIB10 (OP961723). The sequencing reads for recombinant phages underlying this article are available in BioProjects with accession PRJNA488469 [for BIBs 1-4] and PRJNA906721 [for BIBs 6-10].

## SUPPLEMENTARY DATA

Supplementary Table 1

Supplementary Figure 1

## ACKNOWLEDGEMENT

A portion of the work described here was performed in a Science Education Alliance Phage Hunters Advancing Genomics and Evolutionary Science (SEA-PHAGES) course at Lehigh University by J.K. with support provided by the Department of Biological Sciences and the Science Education Alliance of the Howard Hughes Medical Institute (Chevy Chase, MD). We thank SEA-PHAGES team members Daniel Russell and Rebecca Garlena for performing genome sequencing, genome assembly, and deposition of raw reads into the Sequence Read Archive. We thank the Graham Hatfull laboratory for the Island3 phage lysate, and Dr. Michael Kuchka and the phage community for thoughtful feedback about this work.

## TABLES AND FIGURE LEGENDS

**Figure S1.**
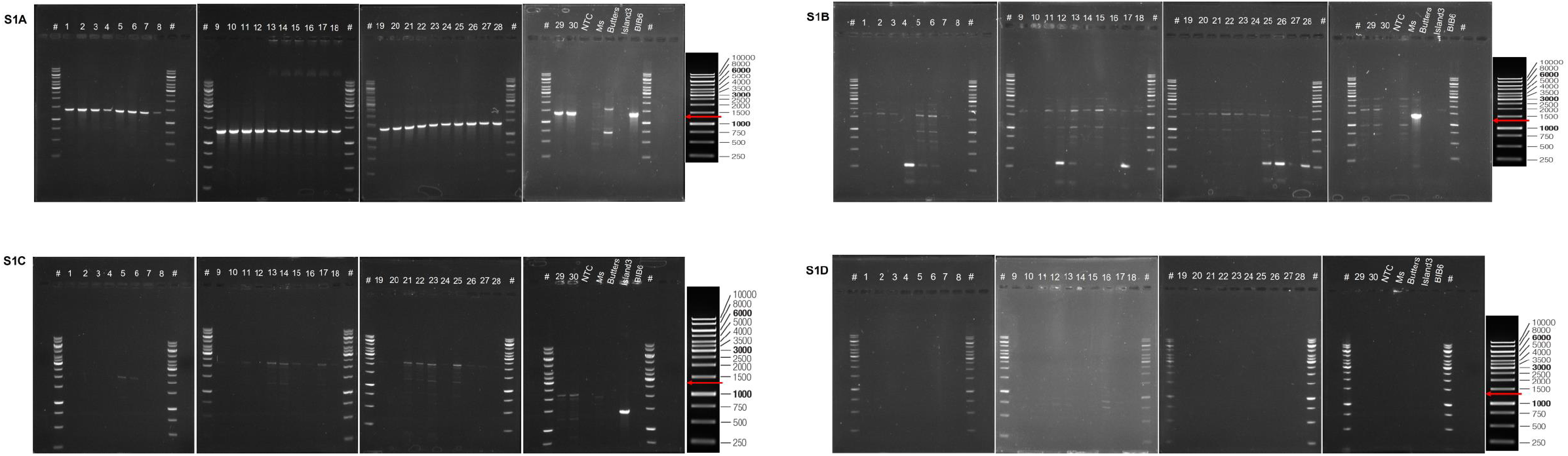
PCR screening of secondary plaques obtained by replating primary plaques (related to Figure 4A). Each secondary plaque was picked into 100 μl of phage buffer and 1 μl used as template for PCR. The first three panels contain plaques 1-28. The fourth panel contains ladder (#), plaques 29, 30, No template control (NTC), *M. smegmatis* mc2155 *(Ms)*, Butters lysate, Island3 lysate, BIB6 lysate, and ladder (#). **A**. PCR performed using Butters-specific primers. All plaques (1-30) showed the expected band (1361bp) as seen in the Butters lysate control lane. **B**. PCR performed using Island3-specific primers. **C**. PCR performed using BIB-specific primers **D**. PCR performed using IBI-specific primers

**Table S1:**
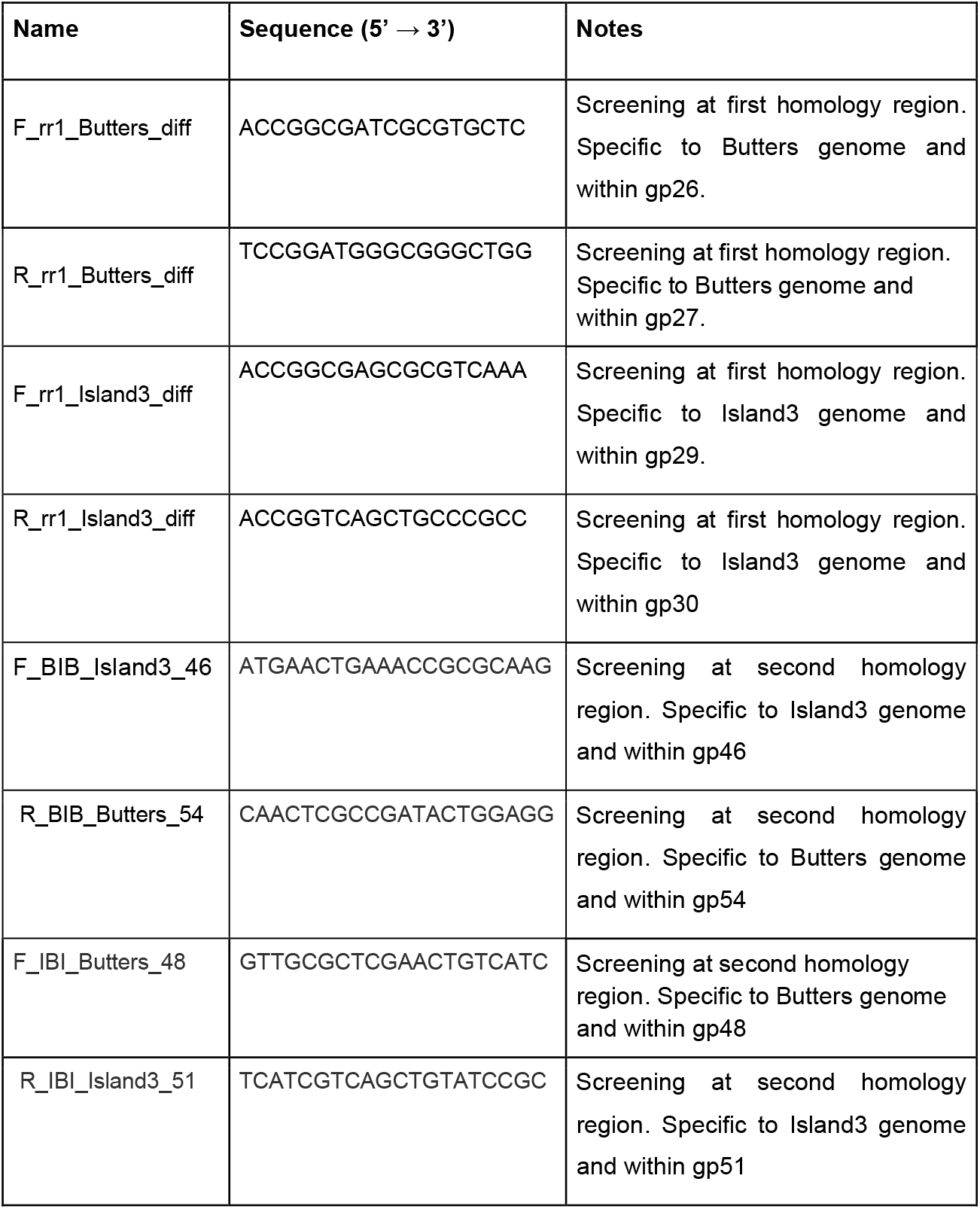
List of primers used for Polymerase Chain Reactions

## REFERENCES

Abedon, S. T. Lysis from without. Bacteriophage 1: 46–49.

Aslam, S., A. M. Courtwright, C. Koval, S. M. Lehman, S. Morales et al., 2019 Early clinical experience of bacteriophage therapy in 3 lung transplant recipients. Am. J. Transplant. 19:.

Bondy-Denomy, J., J. Qian, E. R. Westra, A. Buckling, D. S. Guttman et al., 2016 Prophages mediate defense against phage infection through diverse mechanisms. ISME J. 10: 2854–2866.

Botstein, D. A THEORY OF MODULAR EVOLUTION FOR BACTERIOPHAGES *.:

Broussard, G. W., L. M. Oldfield, V. M. Villanueva, B. L. Lunt, E. E. Shine et al., 2013 Integration-Dependent Bacteriophage Immunity Provides Insights into the Evolution of Genetic Switches. Mol. Cell 49: 237–248.

Caratenuto, R. A., G. O. Ciabattoni, N. J. DesGranges, C. L. Drost, L. Gao et al., 2019 Genome sequences of six cluster n mycobacteriophages, kevin1, nenae, parmesanjohn, shrimpfriedegg, smurph, and spongebob, isolated on mycobacterium smegmatis mc2155. Microbiol. Resour. Announc. 8:.

Casjens, S., 2003 Prophages and bacterial genomics: What have we learned so far? Mol. Microbiol. 49: 277–300.

Clark, A. J., W. Inwood, T. Cloutier, and T. S. Dhillon, 2001 Nucleotide sequence of coliphage HK620 and the evolution of lambdoid phages. J. Mol. Biol. 311: 657–679.

Cresawn, S. G., M. Bogel, N. Day, D. Jacobs-Sera, R. W. Hendrix et al., 2011 Phamerator: A bioinformatic tool for comparative bacteriophage genomics. BMC Bioinformatics 12:.

Dedrick, R. M., C. A. Guerrero-Bustamante, R. A. Garlena, D. A. Russell, K. Ford et al., 2019 Engineered bacteriophages for treatment of a patient with a disseminated drug-resistant Mycobacterium abscessus. Nat. Med. 25: 730–733.

Dedrick, R. M., D. Jacobs-Sera, C. A. Guerrero Bustamante, R. A. Garlena, T. N. Mavrich et al., 2017 Prophage-mediated defence against viral attack and viral counter-defence. Nat. Microbiol. 2:.

Dedrick, R. M., B. E. Smith, R. A. Garlena, D. A. Russell, H. G. Aull et al., 2021 Mycobacterium abscessus strain morphotype determines phage susceptibility, the repertoire of therapeutically useful phages, and phage resistance. MBio 12:.

Delbrt3ck, M. THE GROWTH OF BACTERIOPHAGE AND LYSIS OF THE HOST Introduction and Statement of Main Result.:

Dragoš, A., B. Priyadarshini, Z. Hasan, M. L. Strube, P. J. Kempen et al., 2021 Pervasive prophage recombination occurs during evolution of spore-forming Bacilli. ISME J. 15:.

Guirouilh-Barbat, J., S. Lambert, P. Bertrand, and B. S. Lopez, 2014 Is homologous recombination really an error-free process? Front. Genet. 5:.

Hampton, H. G., B. N. J. Watson, and P. C. Fineran, 2020 The arms race between bacteria and their phage foes. Nature 577: 327–336.

Hatfull, G. F., 2020 Actinobacteriophages: Genomics, Dynamics, and Applications. Annu. Rev. Virol. 7:.

Hendrix, R. W., J. G. Lawrence, G. F. Hatfull, and S. Casjens, 2000 The origins and ongoing evolution of viruses. Trends Microbiol. 8:.

Juhala, R. J., M. E. Ford, R. L. Duda, A. Youlton, G. F. Hatfull et al., 2000 Genomic sequences of bacteriophages HK97 and HK022: Pervasive genetic mosaicism in the lambdoid bacteriophages. J. Mol. Biol. 299:.

Katsura, I., and R. W. Hendrix, 1984 Length determination in bacteriophage lambda tails. Cell 39:.

Kauffman, K. M., W. K. Chang, J. M. Brown, F. A. Hussain, J. Y. Yang et al. Resolving the structure of phage-bacteria interactions in the context of natural diversity.

Kutateladze, M., and R. Adamia, 2008 Phage therapy experience at the Eliava Institute. Med. Mal. Infect. 38:.

Labrie, S. J., J. E. Samson, and S. Moineau, 2010 Bacteriophage resistance mechanisms. Nat. Rev. Microbiol. 8: 317–327.

Law, N., C. Logan, G. Yung, C. L. L. Furr, S. M. Lehman et al., 2019 Successful adjunctive use of bacteriophage therapy for treatment of multidrug-resistant Pseudomonas aeruginosa infection in a cystic fibrosis patient. Infection 47:.

Levine, M., 1957 Mutations in the temperate phage P22 and lysogeny in Salmonella. Virology 3:.

Mageeney, C. M., H. T. Mohammed, M. Dies, S. Anbari, N. Cudkevich et al., 2020 Mycobacterium Phage Butters-Encoded Proteins Contribute to Host Defense against Viral Attack (R. Alegado, Ed.). mSystems 5:.

Martinsohn, J. T., M. Radman, and M. A. Petit, 2008 The λ red proteins promote efficient recombination between diverged sequences: Implications for bacteriophage genome mosaicism. PLoS Genet. 4:.

Mavrich, T. N., and G. F. Hatfull, 2017 Bacteriophage evolution differs by host, lifestyle and genome. Nat. Microbiol. 2:.

Morris, P., L. J. Marinelli, D. Jacobs-Sera, R. W. Hendrix, and G. F. Hatfull, 2008 Genomic characterization of mycobacteriophage giles: Evidence for phage acquisition of host DNA by illegitimate recombination. J. Bacteriol. 190:.

Oliveira, H., M. Sampaio, L. D. R. Melo, O. Dias, W. H. Pope et al., 2019 Staphylococci phages display vast genomic diversity and evolutionary relationships. BMC Genomics 20:.

De Paepe, M., G. Hutinet, O. Son, J. Amarir-Bouhram, S. Schbath et al., 2014 Temperate Phages Acquire DNA from Defective Prophages by Relaxed Homologous Recombination: The Role of Rad52-Like Recombinases. PLoS Genet. 10:.

Pedulla, M. L., M. E. Ford, J. M. Houtz, T. Karthikeyan, C. Wadsworth et al., 2003 Origins of Highly Mosaic Mycobacteriophage Genomes creasingly clear that these phages exert enormous influ-ence over the microbial world (: Wilhelm and Suttle.

Piel, D., M. Bruto, Y. Labreuche, F. Blanquart, S. Lepanse et al., 2021 Genetic determinism of phage-bacteria coevolution in natural populations.

Pires, D. P., H. Oliveira, L. D. R. Melo, S. Sillankorva, and J. Azeredo, 2016 Bacteriophage-encoded depolymerases: their diversity and biotechnological applications. Appl. Microbiol. Biotechnol. 100:.

Pope, W. H., C. A. Bowman, D. A. Russell, D. Jacobs-Sera, D. J. Asai et al., 2015 Whole genome comparison of a large collection of mycobacteriophages reveals a continuum of phage genetic diversity. Elife 4:.

Ramesh, N., L. Archana, M. Madurantakam Royam, P. Manohar, and K. Eniyan, 2019 Effect of various bacteriological media on the plaque morphology of Staphylococcus and Vibrio phages. Access Microbiol. 1:.

Rebekah M. Dedrick, Bailey E. Smith, Madison Cristinziano, Krista G. Freeman, Deborah Jacobs-Sera, Yvonne Belessis, A. Whitney Brown, Keira A. Cohen, Rebecca M. Davidson, David van Duin, Andrew Gainey, Cristina Berastegui Garcia, C.R. Robert George, Ghady, G. F. H., 2022 Phage Therapy of Mycobacterium Infections: Compassionate-use of Phages in Twenty Patients with Drug-Resistant Mycobacterial Disease. Clin. Infect. Dis. 1–6.

Reyes, A., N. P. Semenkovich, K. Whiteson, F. Rohwer, and J. I. Gordon, 2012 Going viral: Next-generation sequencing applied to phage populations in the human gut. Nat. Rev. Microbiol. 10:.

Russell, D. A., and G. F. Hatfull, 2017 PhagesDB: The actinobacteriophage database. Bioinformatics 33: 784–786.

Santoyo, G., and D. Romero, 2005 Gene conversion and concerted evolution in bacterial genomes. FEMS Microbiol. Rev. 29:.

Schneider, C. A., W. S. Rasband, and K. W. Eliceiri, 2012 NIH Image to ImageJ: 25 years of image analysis. Nat. Methods 9:.

Shin, H., J.-H. Lee, H. Yoon, D.-H. Kang, and S. Ryu, 2014 Genomic Investigation of Lysogen Formation and Host Lysis Systems of the Salmonella Temperate Bacteriophage SPN9CC. Appl. Environ. Microbiol. 80:.

The Actinobacteriophage Database | Home.

Touchon, M., A. Bernheim, and E. P. C. Rocha, 2016 Genetic and life-history traits associated with the distribution of prophages in bacteria. ISME J. 10:.

Tzipilevich, E., O. Pollak-Fiyaksel, and S. Ben-Yehuda, 2021 Bacteria elicit a phage tolerance response subsequent to infection of their neighbors. bioRxiv 2021.02.16.428622.

Visconti, N., and M. Delbrück, 1953 THE MECHANISM OF GENETIC RECOMBINATION IN PHAGE. Genetics 38:.

Waterhouse, A. M., J. B. Procter, D. M. A. Martin, M. Clamp, and G. J. Barton, 2009 Sequence analysis Jalview Version 2-a multiple sequence alignment editor and analysis workbench. Bioinforma. Appl. NOTE 25: 1189–1191.

Yahara, K., P. Lehours, and F. F. Vale, 2019 Analysis of genetic recombination and the pan-genome of a highly recombinogenic bacteriophage species. Microb. Genomics 5:.

